# Heart-brain coupling and its relevance for individual trait characteristics

**DOI:** 10.1101/2024.04.30.591871

**Authors:** Suvi Karjalainen, Jan Kujala, Tuija Aro, Tiina Parviainen

## Abstract

A bulk of recent neurophysiological research has focused on how bodily functions are intertwined with neural activity, but the dynamic heart-brain coupling and its relevance for individual trait characteristics remains largely unexamined. Thus, our aim was to investigate how ongoing oscillatory brain activity is modulated by the natural fluctuations in heart rate variability (HRV). We further explored whether heart-brain coupling is associated with individual trait characteristics. Magnetoencephalography (MEG) together with electrocardiography (ECG) were used to record neural activity and HRV during rest. Self-reported trait characteristics were examined using Behavioral Inhibition and Activation Systems Scale (BIS/BAS) and attunement to internal bodily sensations using Body Vigilance Scale (BVS). Statistically significant increases were observed for low HRV vs. high HRV state in alpha and beta power (*p* < 0.05) indicating that oscillatory brain activity is modulated by fluctuations in HRV. Moreover, we demonstrated that heart-brain coupling was associated with self-reported behavioral approach and avoidance tendencies. The results of the moderator analysis further indicated that the relationship between heart-brain coupling and trait characteristics is at least partly moderated by the attunement to internal bodily sensations. Our findings bring insights to the intricate interplay between cardiac and neural signaling and its relationship with individual trait characteristics.

**Highlights:**

- Oscillatory brain activity is modulated by the natural fluctuations in HRV
- Alpha and beta power increase during states of lower parasympathetic activity
- Heart-brain coupling is linked with self-reported individual trait characteristics
- This association is influenced by the attunement to the internal bodily milieu

## Introduction

Human body consists of several systems that are characterized by dynamic rhythms functioning at varying timescales. Neural oscillations, that is rhythmic patterns of brain activity, play a crucial role in the coordination of activity within and across brain areas (Buzsáki and Draguhn, 2004; Thut et al., 2012). Furthermore, neural oscillations are considered as markers of the functional brain states as they are consistently reported to be related to a variety of processes including attention, inhibition, and maintenance of the current cognitive or sensorimotor state (Engel and Fries, 2010; Foxe and Snyder, 2011; Jensen and Mazaheri, 2010; Klimesch, 2012; Klimesch et al., 2007). The spectral characteristics of oscillatory brain activity are partly heritable, but they vary also in state-dependent manner (Pauls et al., 2024; Salmela et al., 2016). Their variance is thus likely to be meaningful for both inter- and intra-individual variability (Haegens et al., 2014). Besides neural networks, bodily systems operate at their characteristic frequency (e.g., heart rate 1.25 Hz). Although bodily rhythms are considerably slower than neural oscillations, they share the basic principles of functional coupling possibly enabling their mutual interaction (Criscuolo et al., 2022; Klimesch, 2018; Nokia and Penttonen, 2022).

Traditionally, the brain has been regarded as the conductor orchestrating the functions of the body, but research evidence is accumulating to show how bodily functions, such as cardiac (Duschek et al., 2015; Thayer et al., 2012), respiratory (Herrero et al., 2018; Karjalainen et al., 2023; Kluger and Gross, 2021), and gastric activity (Richter et al., 2017) influence neural activity in the brain. Furthermore, an extensive network consisting of brainstem, limbic, and sensorimotor regions is influenced by the peripheral physiology (Beissner et al., 2013; Benarroch, 1993; Kluger and Gross, 2021; Rebollo and Tallon-Baudry, 2022; Yang and Feldman, 2018), indicating that the functioning of different bodily systems govern neural activity across cortical and subcortical regions.

The interaction between cardiac and neural activity is especially interesting due to the bidirectional anatomical and functional coupling between the heart and brain (Candia-Rivera et al., 2022; Silvani et al., 2016). Reasonable way to approach heart-brain coupling is via heart rate variability (HRV), the natural fluctuation in the time intervals between consecutive heartbeats. HRV demonstrates the dynamic interplay between sympathetic and parasympathetic nervous systems (Acharya et al., 2006; TFESCNASP Electrophysiology, 1996) providing not only insight into cardiac activity but also a window into the physiological state of the autonomic nervous system. Since HRV reflects an individual’s adaptation to changing internal and external circumstances (Acharya et al., 2006; Shaffer and Ginsberg, 2017), it has been suggested to reflect individual trait characteristics. Indeed, individual differences in HRV have been linked with variability in affective and cognitive processes (Appelhans and Luecken, 2006; Forte et al., 2019; Luque-Casado et al., 2016). Although associations between HRV and the functional properties of neural activity have been observed during rest (Chang et al., 2013; Valenza et al., 2019) and tasks (Lane et al., 2009; Thayer et al., 2012), the dynamic coupling between HRV and ongoing oscillatory brain activity remains an open area for further investigation.

Despite the rather scarce evidence, accumulating findings indicate that the interactions between peripheral physiology and neural activity influence affective and cognitive processes (D’Hondt et al., 2010; Lacey and Lacey, 1978; Parviainen et al., 2022; Varga and Heck, 2017). For instance, the phases of the cardiac cycle influence neural processing of fear and threat (Garfinkel et al., 2014). Similarly, the phases of the respiratory cycle modulate perceptual sensitivity (Kluger et al., 2021) and learning-related processes (Waselius et al., 2019; Waselius et al., 2022). The aforementioned studies provide mechanistic understanding on how the interaction between bodily functions and neural activity may influence processing of external inputs. However, less attention has been paid to individual variation in body-brain interaction and how it may affect the behavioral tendencies to approach and interact with the external world.

Temperamental characteristics represent individually unique, neurobiologically based behavioral tendencies in the way an individual experiences and reacts to the world (Gray, 1991). Innate behavioral approach and avoidance tendencies have been associated with distinctive patterns of neural activity and physiological reactivity (Heponiemi et al., 2004; Kennis et al., 2013; Knyazev et al., 2002). Moreover, temperamental characteristics are reported to be correlated with the synchrony between the central and autonomic nervous system functioning, indicating that body-brain coupling and especially the afferent information flow from the periphery to the brain may contribute to shaping individual trait characteristics (Shokri-Kojori et al., 2018).

However, given the recent evidence (Critchley and Garfinkel, 2018; Garfinkel et al., 2015; Engelen et al., 2023; Tsakiris and Critchley, 2016), it may be that the sensitivity to or accuracy in perceiving internal bodily signals (i.e., interoception), rather than body-brain interaction per se, underlies the relevance of body-brain interaction for subjective experience and behavior. Previous findings have indicated that modulations in cortical responses to cardiac activity are linked with attention to and accuracy in the perception of interoceptive signals (Coll et al., 2021; Petzschner et al., 2019; Pollatos et al., 2005). Furthermore, sensitivity to bodily sensations has been reported to be associated with behavioral avoidance tendency (Lyyra and Parviainen, 2018). Intriguingly, the tendency to attend to information from the internal milieu is associated with modulations in oscillatory brain activity (Hanslmayr et al., 2011; Villena-González et al., 2017). Considering the interplay between bodily functions and neural activity, the attunement to the internal bodily sensations could play a crucial role in determining whether and how body-brain interaction is linked with individual trait characteristics.

To add on to the existing evidence on the coupling between bodily functions and neural activity, we focused on the unexplored interplay between ongoing fluctuations in cardiac and neural signaling. Specifically, we investigated how the spectral characteristics of ongoing oscillatory brain activity are modulated by the natural fluctuations in HRV. Furthermore, since the relevance of body-brain interaction for individual trait characteristics remains largely unexamined, we explored whether individual variation in heart-brain coupling is associated with self-reported trait characteristics, namely behavioral approach and avoidance tendencies, either directly or moderated by the attunement to the internal bodily sensations.

## Materials and methods

### Participants

This study consists of two cohorts of participants who were involved in this study in 2018-2019 (n = 12) and 2020-2021 (n = 28). From the initial 40 participants, seven were excluded due to incomplete questionnaire data (n = 1) or low quality of the HRV and/or MEG data (n = 6) resulting in the final sample of 33 participants (24 female, age 25.39 ± 6.87 years [mean ± SD], range 19-57 years). Exclusion criteria included a history of cardiovascular or respiratory disease, head trauma, intellectual disability, neurological or psychiatric condition, use of medication affecting the nervous system, and claustrophobia. Additionally, participants included in this study had normal or corrected-to-normal hearing and vision. Ethical approval was received from the local Ethics Committee of the University of Jyväskylä and all the study procedures were conducted in accordance with the Declaration of Helsinki. Prior to any study procedures, participants were informed about the study and a written informed consent was obtained from all participants.

### Procedure

The data collection of the first cohort (n = 12, of which 8 participants were included in the final sample) consisted of one visit (3 hours) in the MEG laboratory of Centre for Interdisciplinary Brain Research (CIBR, Jyväskylä, Finland). Participants of the first cohort completed the questionnaires at home after the visit. The data collection of the second cohort (n = 28, of which 25 participants were included in the final sample) consisted of two visits in CIBR. During the first visit (1.5 hours) cognitive tests were conducted and a set of questionnaires was given to the participants to be filled out at home before the MEG laboratory visit (2.5 hours). Visits to the MEG laboratory were scheduled either in the morning (9-12 am) or in the afternoon (12-15 pm) to minimize the effect of circadian rhythm and participants were instructed to refrain from nicotine and caffeine for at least 3 hours prior to the visit.

During the MEG recordings participants were seated upright in the MEG device and their positions were stabilized with pillows. They were also instructed to avoid any excessive movements during the recordings to minimize head movements. Participants were instructed to sit relaxed, keep their eyes on a fixation cross (distance from the participant’s eyes ∼ 2,5 m), and breathe through their nose for the duration of 8 minutes to record neural activity and HRV during rest. The experimental design also included a resting state condition with eyes closed, volitionally controlled deep and square breathing conditions, and a heartbeat discrimination task that are not reported in this paper.

### Data recordings

MEG data was collected using a whole-head 306-sensor (102 magnetometer channels and 204 planar gradiometer channels) Elekta Neuromag TRIUX system (MEGIN Oy, Helsinki, Finland) in a magnetically shielded sound-attenuated room. Data were sampled at 1000 Hz and filtered at 0.1-330 Hz. Head position with respect to the MEG sensors was continuously monitored using five head position indicator (HPI) coils that were attached to the scalp of the participant (three HPI coils placed on the forehead and one behind each ear). Moreover, prior to the MEG recordings the anatomical landmarks (left and right preauricular points, nasion), position of each HPI coil, and an additional set of points (∼ 150) distributed over the scalp were digitized using the Polhemus Isotrak digital tracker system (Polhemus, Colchester, VT, USA) to allow co-registration with the MRI template.

Electro-oculogram (EOG), electrocardiogram (ECG), and respiratory signals were recorded simultaneously with the MEG data. To detect eye blinks and saccades an EOG electrode pair was placed diagonally above the right eye and below the left eye. ECG electrodes and a ground electrode were attached below each clavicle and on the right clavicle, respectively. Respiration was monitored with a respiratory belt placed around the participant’s lower chest. During the data collection of the first cohort of participants a piezo-based breathing sensor (Spes Medica, Genova, Italy) was used, whereas an in-house developed strain gauge breathing sensor was used for the second cohort of participants.

### Preprocessing and analysis of cardiac data

ECG data from each participant was extracted from the original data as a comma-separated values file. NeuroKit2, an open-source Python toolbox for neurophysiological signal processing (Makowski et al., 2021), was used for preprocessing and extracting HRV indices from the ECG data. Preprocessing of the ECG signals involved minimizing noise, artifact removal, and R-peak detection. The following well-established cardiac measures across the whole resting state recording (8 min) were computed from the preprocessed ECG signal: mean heart rate (HR mean), mean inter-beat-interval (IBI mean), and the root mean square of the differences of successive RR intervals (RMSSD). To define the low and high HRV states based on the cardiac activity, the preprocessed ECG signal was segmented into 16 consecutive 30 second time windows and RMSSD, a time-domain measure of HRV reflecting parasympathetic activity (Bertsch et al., 2012), was calculated for each segment. Then, the segments in which RMSSD was at its highest (i.e. high HRV) and lowest (i.e., low HRV) were identified. To increase the contrast between the low and high HRV states, 6 segments instead of 8 segments per category were included in the further analyses.

### Preprocessing and analysis of MEG data

MaxFilter™ 3.0 software (MEGIN, Helsinki, Finland) was applied to remove external magnetic interference from the MEG data using the temporal extension of the signal-space separation (tSSS) method (Taulu et al., 2004; Taulu and Kajola, 2005; Taulu and Simola, 2006). Furthermore, the head position was estimated for head movement compensation and the median head position across all participants’ MEG recordings was used as a reference head position. MEG data was downsampled resulting in a sampling rate of 500 Hz. Thereafter MEG data was exported to Meggie, an MNE-python-based graphical user interface (Heinilä and Parviainen, 2022), where independent component analysis (ICA) was used to extract, visually inspect, and manually remove common artifacts of cardiac and ocular origin. Since gradiometers are optimal for recording data from superficial sources and they have a narrow spatial sensitivity pattern, the subsequent analyses were conducted using only planar gradiometers.

To explore whether and how the functional state of the brain differs between low vs. high HRV state, MEG data was first segmented into 16 consecutive 30 second time windows matching the timing of the HRV segmentation. Then the windows in which RMSSD was at its highest (i.e., high HRV) or lowest (i.e., low HRV) were combined and included in further analysis. For both high and low HRV, 6 time windows were combined, yielding 3 minutes of MEG data per category. Next, cross spectral density (CSD) matrices were calculated for both low and high HRV states within the frequency range of 2 – 45 Hz (length of the window 2048, 50 % overlap, Hanning window).

The spatial distribution of modulations in oscillatory brain activity was estimated using Dynamic Imaging of Coherent Sources (DICS; Gross et al., 2001). Since the DICS analysis was conducted in a template brain (FreeSurfer fsaverage-5.1.0 template; Fischl, 2012), the forward model of one participant utilizing a single-compartment realistic boundary element model was used to define the distribution of points across the parcels for all participants. A customized version of the Destrieux anatomical parcellation (Destrieux et al., 2010) with 69 parcels per hemisphere that was constructed using a merge-and-split approach to produce uniform-sized parcels (Ala-Salomäki et al., 2021) was used as the parcellation. The frequency-domain beamformer was implemented using a common weights approach, in which the weights were defined by combining both CSD matrices (i.e., low and high HRV states). Beamformer estimates at the parcel-level across the whole brain were calculated for all participants. Here, joint beamformer weights were estimated across the high and low HRV states across the whole 2– 45 Hz band for each parcel. These weights were then applied separately to the two states and each frequency bin, yielding parcel-level spectra for both states. Finally, Fitting oscillations and one over f (FOOOF) algorithm (Donoghue et al., 2020) was applied to extract both periodic (i.e., power) and aperiodic components (i.e., exponent and offset) of the power spectra in each parcel. For periodic activity, the power within each frequency band of interest (theta, 4 – 7 Hz; alpha, 8 – 13 Hz; beta, 15 – 25 Hz; gamma 35 – 45 Hz) was calculated by subtracting the aperiodic activity from the original spectra.

### Individual trait characteristics

BIS/BAS scale, a 20-item self-report questionnaire, was applied to assess the responsiveness of the two motivational systems, the behavioral inhibition system (BIS) and approach system (BAS) (Carver and White, 1994). BIS is considered to reflect the sensitivity to negative affect, punishment, and non-reward leading to behaviors such as avoidance and inhibition to avoid negative outcomes, whereas BAS is associated with positive affect, non-punishment, and reward resulting in goal-directed behaviors (Carver and White, 1994; Gray, 1991). BIS/BAS scale is divided into four subscales (BIS, BAS drive, BAS fun seeking, and BAS reward responsiveness) each showing good reliability (Carver and White, 1994). Each item is answered on a 5-point Likert scale ranging from 1 (not at all like me) to 5 (extremely like me). To test the reliability of the BIS/BAS scale in our sample, the internal consistency was assessed using Cronbach’s alpha coefficient. In the present study, internal consistency yielded a Cronbach’s *α* of 0.81 for both BIS and BAS drive, 0.70 for BAS fun seeking, and 0.51 for BAS reward responsiveness subscales.

### Attunement to internal bodily sensations

Body Vigilance Scale (BVS), a four-item self-report scale, was used to assess the tendency to attend to internal bodily sensations (Schmidt et al., 1997). The first two items assess the degree of attentional focus and perceived sensitivity to changes in bodily sensations on an 11-point scale ranging from 0 (not at all like me) to 10 (extremely like me). Third, the average amount of time spent attending to bodily sensations is assessed using an 11-point scale ranging from 0 (no time) to 100 (all of the time). When calculating the total BVS score, the score of the third item is divided by 10 to match the scales of the other items. The fourth item measures the degree of attention to 15 different anxiety-related somatic sensations on a scale from 0 (none) to 10 (extreme). The ratings of the 15 sensations are averaged to yield the overall score for the fourth item. The total BVS score is the sum of the scores across items ranging from 0 to 40. BVS has been reported to be a unidimensional scale with good internal consistency and adequate test-retest reliability (Schmidt et al., 1997). The internal consistency yielded a Cronbach’s *α* of 0.77 for the total BVS score.

### Statistical analyses

Statistical analyses were performed using IBM SPSS Statistics 28.0 (IBM Corp., Armonk, New York, USA) and MATLAB (R2020b, The MathWorks Inc., Natick, MA, USA). To assess the differences in cardiac activity between low vs. high HRV state, the average RMSSD across the segments where RMSSD is at its lowest (RMSSD low) and highest (RMSSD high) were calculated using SPSS. According to the Kolmogorov-Smirnov test RMSSD low (*D*(33) = 0.189, *p* = 0.004) and RMSSD high (*D*(33) = 0.226, *p* < 0.001) variables were not normally distributed. Thus, a non-parametric Wilcoxon Signed-Rank test was performed to investigate the differences in RMSSD between low vs. high HRV state.

The differences between low vs. high HRV states in periodic (i.e., power) and aperiodic (i.e., exponent and offset) activity on the parcel-level were tested by dependent samples t-test using MATLAB. The *p*-values across parcels were corrected for multiple comparisons using false discovery rate (FDR). Next, we investigated the associations between heart-brain coupling measures and self-reported trait characteristics. For this, we first calculated the differences in periodic and aperiodic activity between low vs. high HRV states (HRV_diff_) for each parcel in which statistically significant effects (*p* < 0.05, corrected for multiple comparisons) were observed. Thereafter, Pearson correlation coefficients were computed to examine the associations between HRV_diff_ and BVS, BIS, BAS drive, BAS fun seeking, and BAS reward responsiveness variables.

Finally, we tested whether heart-brain coupling (quantified as HRV_diff_) is directly related to the behavioral approach and avoidance tendencies or whether this association is moderated by the attunement to the internal bodily sensations. For this purpose, a multiple linear regression model in SPSS was constructed that included measures of HRV_diff_, BVS, and their interaction HRV_diff_ x BVS as independent variables and BIS/BAS as the dependent variable. All variables were standardized and entered simultaneously to the model. The interaction variable was computed by multiplying the standardized HRV_diff_ with the standardized BVS score. In case a significant interaction effect was observed, simple slopes analysis (online tool by Preacher et al., 2004) was applied to assess the relationship between heart-brain coupling (HRV_diff_) and behavioral tendency (BIS/BAS) at high (+1 SD) or low (−1 SD) levels of the moderator (BVS). The histogram and the normal P-P plot of standardized residuals indicated that the errors were approximately normally distributed. Moreover, the scatterplot of standardized residuals showed that the assumptions of homoscedasticity (i.e., homogeneity of variance) and linearity were met. Since all variance inflation factors (VIF) were < 1.3, multicollinearity was not a concern.

## Results

### Descriptive statistics

Descriptive statistics of cardiac metrics and self-reported trait characteristics are reported in Table 1. Measures of cardiac activity across the whole resting state recording, that is HR mean (67.66 ± 8.21 beats/min), IBI mean (901.15 ± 118.66 ms), and RMSSD (64.36 ± 46.62 ms), fell within the normal reference range for adults (Nunan et al., 2010; Shaffer and Ginsberg, 2017). Expectedly, statistically significant difference was observed in mean RMSSD between low and high HRV states (Z = −5.012, p < 0.001).

**Table 1.** Descriptive statistics of HRV metrics and self-report questionnaire scores.

|  | Mean | SD |
| --- | --- | --- |
| **HR mean total** | 67.66 | 8.21 |
| **IBI mean total** | 901.15 | 118.66 |
| **RMSSD total** | 64.36 | 46.62 |
| **RMSSD low** | 57.11 | 41.66 |
| **RMSSD high** | 75.20 | 54.33 |
| **BVS** | 17.76 | 7.31 |
| **BAS drive** | 14.48 | 2.98 |
| **BAS fun seeking** | 14.76 | 3.23 |
| **BAS reward responsiveness** | 19.36 | 2.76 |
| **BIS** | 23.94 | 5.50 |

*Abbreviations: HR, heart rate; IBI, inter-beat-interval; RMSSD, the root-mean square of the differences of successive RR intervals; BVS, Body Vigilance Scale; BAS, Behavioral Activation Scale; BIS, Behavioral Inhibition Scale; SD, standard deviation*

### Periodic and aperiodic activity in low vs. high HRV states

To investigate whether the functional state of the brain was modulated by the natural fluctuations in HRV, we explored the spatial distribution of differences in periodic (theta, alpha, beta, and gamma power) and aperiodic (exponent and offset) activity during low vs. high HRV states. The parcel-level dependent samples t-test results (*p* < 0.05, corrected for multiple comparisons) demonstrated an increase in alpha and beta power during low HRV state in comparison with high HRV state (Fig. 1, see Appendix for uncorrected results). Statistically significant differences in alpha power were detected in the left posterior superior temporal cortex and left insular cortex. In beta power, statistically significant effects were observed in the left anterior and posterior temporal cortex, left dorsolateral prefrontal cortex, left posterior middle frontal cortex, left middle occipital cortex, and left medial occipitotemporal cortex. Aperiodic activity did not differ between low vs. high HRV states. Altogether, these findings demonstrate that the spectral characteristics of the ongoing brain oscillatory activity were modulated by the fluctuations in HRV.

**Fig. 1.**
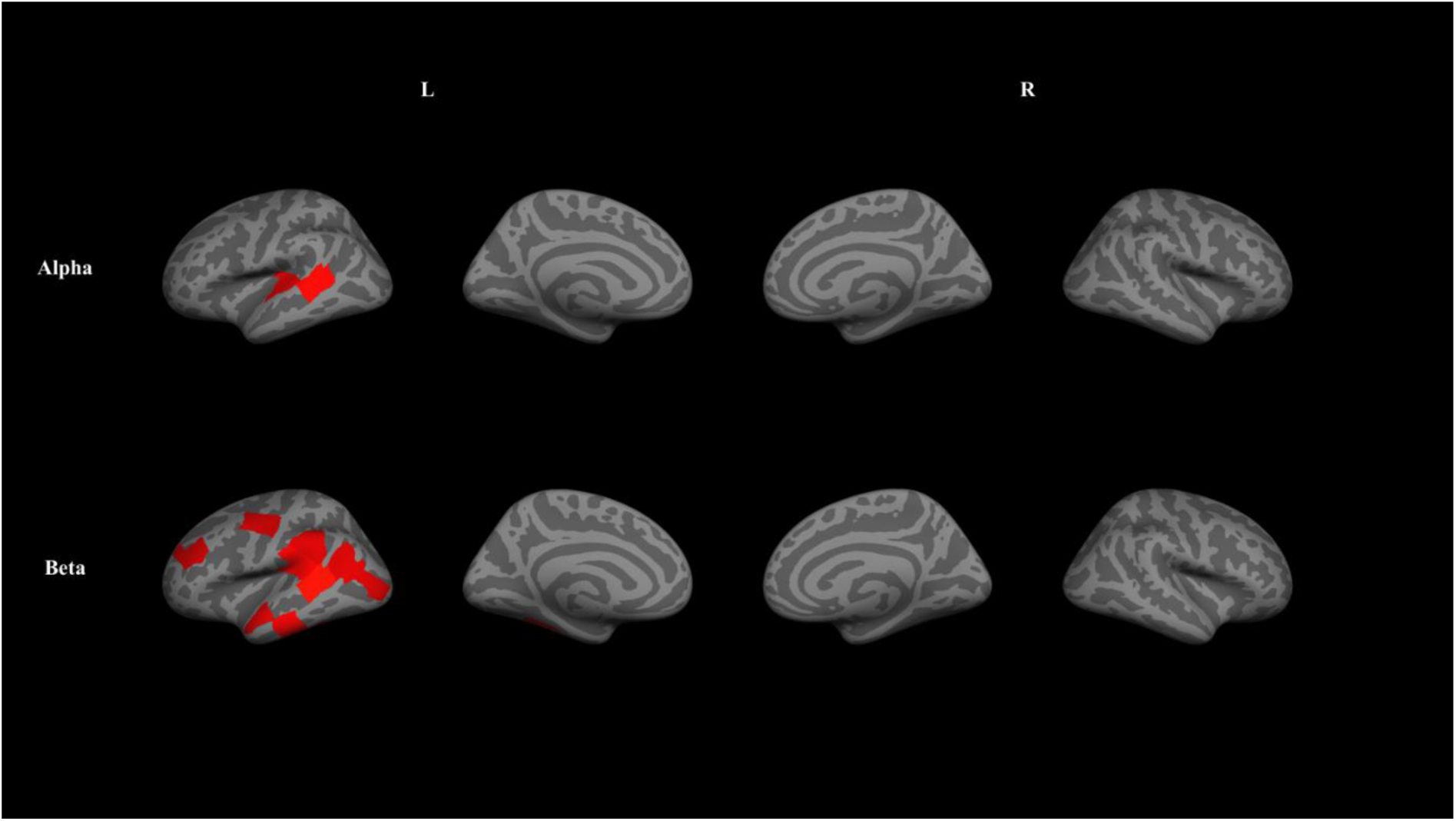
Modulations of periodic activity between low vs. high HRV states. Statistically significant increases (*p* < 0.05, corrected for multiple comparisons) were observed in alpha and beta power between low vs. high HRV states. Red color indicates statistically significant increases in alpha and beta power between low vs. high HRV states.

### Correlations between heart-brain coupling and individual trait characteristics

Correlations between heart-brain coupling (HRV_diff_) and self-reported trait characteristics were investigated in alpha (α) and beta (β) frequency bands in which statistically significant modulations were observed between low vs. high HRV states (see Table 2). Statistically significant correlations were observed between HRV_diff_ measures and between BIS/BAS subscales. The results regarding associations between HRV_diff_ and BIS/BAS demonstrated negative correlation between BIS and HRV_diff_ α in the left temporoparietal cluster (covering left supramarginal gyrus, planum temporale, anterior transverse temporal gyrus, transverse temporal sulcus, and posterior segment of the lateral sulcus) (*r* = −0.352, *p* = 0.044). This correlation signifies that the larger the difference in the alpha power in the low vs. high HRV states was, the weaker was the behavioral avoidance tendency.

**Table 2.** Pearson correlation coefficients between measures of heart-brain coupling and self-reported trait characteristics.

| | Temp<br>par1 $\alpha$ | Sup<br>temp3 $\alpha$ | Mid<br>front $\beta$ | Occ<br>temp $\beta$ | Occ $\beta$ | Temp<br>par2 $\beta$ | Temp<br>par3 $\beta$ | Precent $\beta$ | Sup<br>temp1 $\beta$ | Sup<br>temp3 $\beta$ | Sup<br>temp4 $\beta$ | Inf<br>temp $\beta$ | BVS | BAS drive | BAS fun. | BAS<br>reward | BIS |
| --- | --- | --- | --- | --- | --- | --- | --- | --- | --- | --- | --- | --- | --- | --- | --- | --- | --- |
| Temppar1 $\alpha$ | | | | | | | | | | | | | | | | | |
| Suptemp3 $\alpha$ | **0.428\*** | | | | | | | | | | | | | | | | |
| Midfront $\beta$ | 0.174 | 0.145 | | | | | | | | | | | | | | | |
| Occtemp $\beta$ | **0.420\*** | 0.095 | 0.261 | | | | | | | | | | | | | | |
| Occ $\beta$ | **0.415\*** | 0.266 | **0.402\*** | **0.631\*\*** | | | | | | | | | | | | | |
| Temppar2 $\beta$ | 0.203 | 0.166 | -0.024 | **0.489\*\*** | 0.170 | | | | | | | | | | | | |
| Temppar3 $\beta$ | 0.066 | 0.030 | -0.108 | **0.397\*** | 0.304 | **0.474\*\*** | | | | | | | | | | | |
| Precent $\beta$ | 0.274 | 0.169 | **0.415\*** | -0.149 | 0.049 | 0.152 | -0.069 | | | | | | | | | | |
| Suptemp1 $\beta$ | 0.239 | 0.232 | 0.236 | 0.183 | 0.056 | **0.372\*** | 0.161 | 0.217 | | | | | | | | | |
| Suptemp3 $\beta$ | 0.190 | 0.275 | **0.562\*\*** | **0.574\*\*** | **0.534\*\*** | **0.456\*\*** | 0.253 | 0.336 | **0.395\*** | | | | | | | | |
| Suptemp4 $\beta$ | **0.344\*** | 0.105 | 0.221 | **0.605\*\*** | **0.825\*\*** | 0.296 | **0.647\*\*** | 0.026 | 0.064 | **0.451\*\*** | | | | | | | |
| Inftemp $\beta$ | 0.287 | 0.329 | **0.528\*\*** | **0.378\*** | 0.272 | 0.189 | 0.177 | 0.271 | **0.571\*\*** | **0.559\*\*** | 0.215 | | | | | | |
| BVS | -0.192 | -0.149 | -0.152 | -0.185 | -0.081 | 0.105 | 0.109 | -0.006 | -0.086 | -0.015 | -0.008 | -0.288 |  |  |  |  |  |
| BAS drive | 0.003 | -0.060 | 0.096 | 0.146 | **0.414\*** | 0.100 | 0.238 | 0.034 | -0.264 | 0.093 | 0.343 | 0.089 | -0.030 |  |  |  |  |
| BAS fun seeking | 0.152 | 0.074 | 0.079 | 0.322 | 0.168 | 0.053 | 0.051 | 0.045 | 0.139 | 0.012 | 0.101 | **0.358\*** | **-0.500\*\*** | 0.090 |  |  |  |
| BAS reward | -0.199 | 0.052 | -0.079 | -0.065 | 0.088 | 0.056 | 0.108 | 0.107 | -0.096 | 0.075 | 0.048 | 0.063 | -0.057 | **0.464\*\*** | **0.347\*** |  |  |
| BIS | **-0.352\*** | -0.305 | -0.312 | -0.228 | -0.233 | -0.050 | -0.052 | -0.313 | -0.141 | -0.135 | -0.235 | -0.259 | **0.627\*\*** | -0.021 | -0.275 | 0.160 |  |

Abbreviations: *par*, parietal; *precent*, precentral; *occ*, occipital; *temp*, temporal; *inf*, inferior; *mid*, middle; *sup*, superior; 1-4, subdivisions 1-4; $\alpha$ , alpha; $\beta$ , beta; *BVS*, Body Vigilance Scale; *BAS*, Behavioral Activation Scale; *BIS*, Behavioral Inhibition Scale; \* $p < 0.05$ , \*\* $p < 0.01$

Moreover, positive correlations were found between BAS drive and HRV_diff_ β in the left occipital cluster (covering left middle occipital gyrus, middle occipital sulcus, lunatus sulcus, superior occipital gyrus, superior occipital sulcus, and transverse occipital sulcus) (*r* = 0.414, *p* = 0.017) as well as BAS fun seeking and HRV_diff_ β in the left temporal cluster (covering left inferior temporal gyrus, inferior temporal sulcus, and middle temporal gyrus) (*r* = 0.358, *p* = 0.041). These findings demonstrate that the larger the difference in the beta power in the low vs. high HRV state was, the stronger were the behavioral approach tendencies. HRV_diff_ measures were not correlated with BAS reward responsiveness and BVS. To summarize, these findings indicated that heart-brain coupling was associated with behavioral approach and avoidance tendencies, but not with the attunement to the internal bodily sensations (see Fig. 2).

**Fig. 2.**
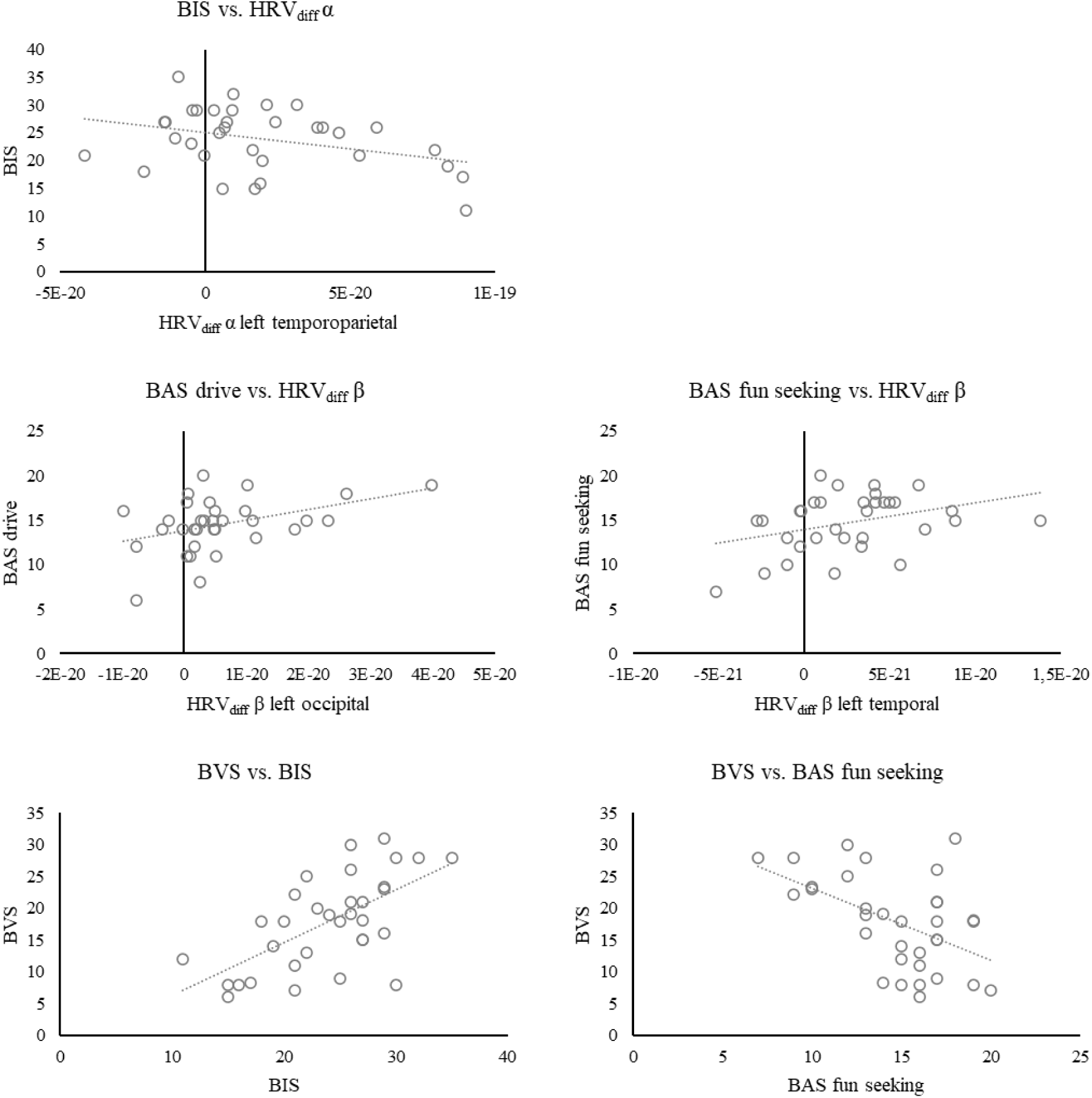
Associations between heart-brain coupling and individual trait characteristics. Scatterplots depicting associations between measures of heart-brain interaction and self-reported individual trait characteristics showing statistically significant correlations (p < 0.05). *Abbreviations: HRV_diff_, difference in the power of neural activity between low vs. high HRV state; α, alpha; β, beta; BVS, Body Vigilance Scale; BAS, Behavioral Activation Scale; BIS, Behavioral Inhibition Scale*

When interactions between different self-reported trait characteristics were examined, positive correlation was observed between BVS and BIS (*r* = 0.627, *p* < 0.001), whereas BVS was negatively correlated with BAS fun seeking (*r* = −0.500, *p* = 0.003). That is, higher levels of attunement to the internal bodily sensations were related to stronger behavioral avoidance tendencies and to weaker behavioral approach tendencies (see Fig. 2). The results thus imply that individuals with heightened attunement to the internal milieu tend to avoid negative outcomes and are less prone to seek novel experiences and excitement. While behavioral approach and avoidance tendencies showed opposite associations with the attunement to internal bodily sensations and were also inversely but not significantly associated with each other (*r* = −0.275, *p* = 0.121), they should not be considered as the opposite ends of the same spectrum but separate, complementary systems that serve distinct functions in regulating behavior (Carver and White, 1994; Jorm et al., 1998).

### Moderator analysis

Since statistically significant correlations were observed between HRV_diff_ α and BIS, HRV_diff_ β and BAS fun seeking as well as between BVS and both BIS and BAS fun seeking variables, we tested whether heart-brain coupling had an independent contribution on behavioral tendencies or whether it was moderated by the attunement to internal bodily sensations.

The results of the multiple linear regression analyses are summarized in Fig. 3. Together HRV_diff_ α, BVS, and HRV_diff_ α x BVS explained 39.3 % (adjusted *R^2^* = 0.393) of the variance in BIS. The results indicated that HRV_diff_ α was not statistically significantly related to BIS (*p* = 0.097). BVS, in turn, was directly related to BIS (*B* = 0.584, *t* = 4.131, *p* < 0.001), but did not moderate the relationship between HRV_diff_ α and BIS (*p* = 0.866).

**Fig. 3.**
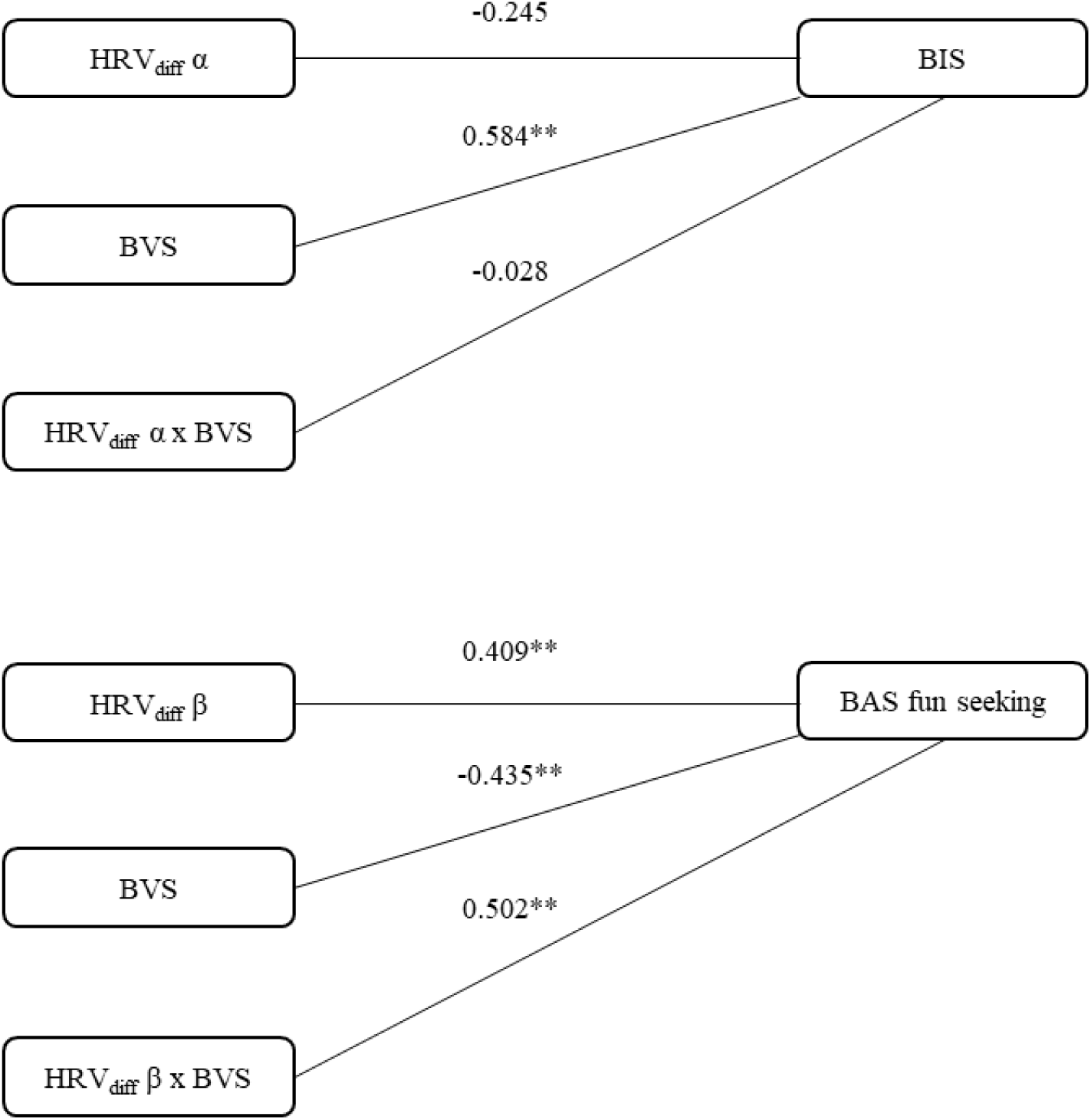
Interaction between heart-brain coupling and individual trait characteristics moderated by the attunement to internal bodily sensations. Statistical diagrams of moderation analyses with unstandardized regression coefficients, where the interaction between heart-brain coupling and behavioral tendencies is moderated by the attunement to internal bodily sensations. *Abbreviations: HRV_diff_, difference in the power of neural activity between low vs. high HRV state; α, alpha; β, beta; BVS, Body Vigilance Scale; BAS, Behavioral Activation Scale; BIS, Behavioral Inhibition Scale; * p < 0.05, ** p < 0.01*

Together HRV_diff_ β, BVS, and HRV_diff_ β x BVS explained 49.5 % (adjusted *R^2^* = 0.495) of the variance in BAS fun seeking. The results demonstrated that HRV_diff_ β was directly related to BAS fun seeking (*B* = 0.409, *t* = 2.951, *p* = 0.006). Similarly, BVS was directly related to BAS fun seeking (*B* = −0.435, *t* = −3.320, *p* = 0.002) and it also moderated the relationship between HRV_diff_ β and BAS fun seeking (*B* = 0.502, *t* = 3.924, *p* < 0.001).

The simple slopes analysis (see Fig. 4) revealed that the positive relationship between HRV_diff_ β and BAS fun seeking was statistically significant among individuals with high BVS (+1 SD) (*B* = 0.911, *p* < 0.001), but not among individuals with low BVS (−1 SD) (*B* = −0.093, *p* = 0.545). To sum up, these results jointly indicate that the relationship between heart-brain coupling and behavioral tendencies is at least partly moderated by the attunement to internal bodily sensations.

**Fig. 4.**
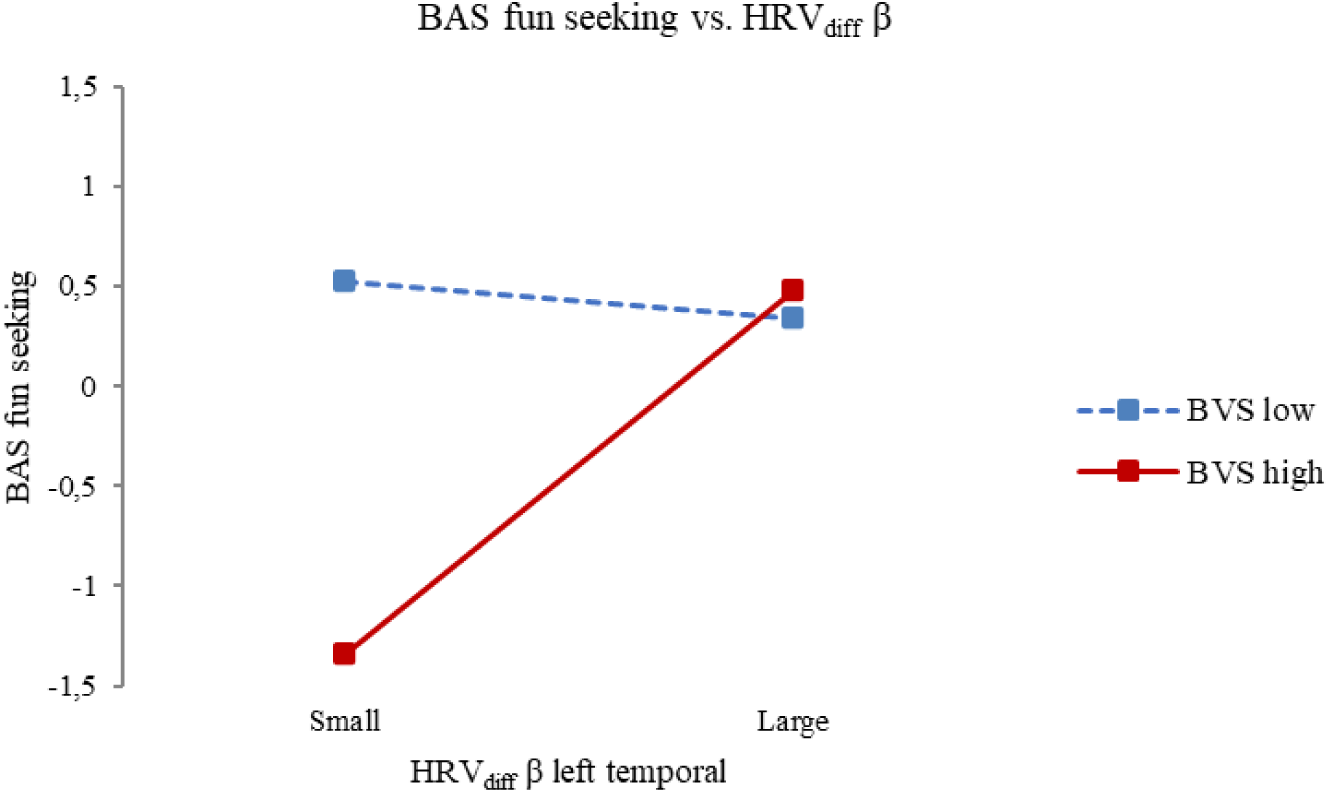
The attunement to internal bodily sensations as a moderator in the association between heart-brain coupling and behavioral approach tendency. The entries “Low” and “High” refer to −1 and +1 standard deviation below and above the sample mean. *Abbreviations: HRV_diff_, difference in the power of neural activity between low vs. high HRV state; β, beta; BAS, Behavioral Activation Scale; BVS, Body Vigilance Scale*

## Discussion

The purpose of this study was to investigate heart-brain coupling and its association with individual trait characteristics. First, we determined whether and how the ongoing oscillatory brain activity is modulated by the fluctuations in HRV, a measure of the functional state of the autonomic nervous system. Increased alpha and beta power were observed during low HRV state in comparison with high HRV state indicating that neural activity responds to the dynamic fluctuations in parasympathetic activity. Second, we examined the relationship between heart-brain coupling and self-reported individual trait characteristics. We demonstrated that individual variation in heart-brain coupling was associated with behavioral approach and avoidance tendencies. Furthermore, our findings indicated that this association was at least partly moderated by the attunement to the internal bodily milieu. Taken together, our results suggest that heart-brain coupling is linked with individual trait characteristics and this relationship is influenced by how we attend to our internal bodily sensations.

Our findings indicated that the spectral characteristics of the ongoing oscillatory brain activity are modulated by the natural fluctuations in HRV during rest. Specifically, an increase in alpha and beta power was observed during states of lower parasympathetic activity as reflected by lower variability in beat-to-beat intervals in comparison with states of higher parasympathetic activity. Our results are in line with and extend earlier studies on heart-brain coupling. In accordance with the current results, a recent study demonstrated that the dynamics of cardiac sympathetic-vagal activity influence large-scale neural network organization in alpha, beta, and gamma frequency bands (Candia-Rivera et al., 2024). Similarly, another study observed robust coupling between HRV and the amplitude of neural oscillations in several frequency bands (Sargent et al., 2024). Based on these findings it seems that the afferent visceral signals may shape the functioning of neural networks. Furthermore, our results are, in general, in accordance with earlier studies demonstrating that parasympathetic activity is associated with the power in the alpha and beta frequency bands (Chang et al., 2011; Takahashi et al., 2005; but see Magosso et al., 2019). However, in earlier studies the relationship between cardiac and neural activity has been approached at coarse level in terms of temporal dynamics (e.g., across tasks or states) and with a limited spatial scope (e.g., specific regions of interest). Our approach, in turn, focused on investigating the dynamic fluctuations in heart-brain coupling across the whole brain at the source-level within a resting state condition that is typically characterized by the dominance of parasympathetic activity. Therefore, our results corroborate earlier evidence showing that neural activity is intertwined with cardiac activity and provide further confirmation that fluctuations in parasympathetic activity modulate the ongoing oscillatory brain activity.

While the current study demonstrated that the ongoing oscillatory brain activity is modulated by dynamic fluctuations in parasympathetic activity, the functional relevance of these findings remains undetermined. However, the existing evidence on the functional role of oscillatory brain activity may shed light on the relevance of heart-brain coupling. Namely, an inverse association between the power in alpha and beta frequency bands and neural excitability has been widely documented (Chapeton et al., 2019; Iemi et al., 2022; Waldhauser et al., 2012). Additionally, modulations in the ongoing oscillatory brain activity have been linked with fluctuations between states in which the attention is directed either towards internal or external milieu (Hanslmayr et al., 2011; Knyazev et al., 2011). If we interpret our findings within this framework, the results indicate that the naturally occurring fluctuations in neural activity may reflect inherent fluctuations in neural excitability and allocation of attentional resources that are possibly driven by subtle changes in the state of the autonomic nervous system. Specifically, lower parasympathetic activity associated with higher alpha and beta power could reflect lower neural excitability to external input and allocation of attentional resources towards the internal bodily milieu.

To understand the functional relevance of our findings regarding heart-brain coupling, it is essential to recognize the bidirectional nature of this interaction. The causal evidence concerning the directionality of cardiac and neural signaling primarily relies on invasive animal studies that have demonstrated the afferent and efferent pathways connecting heart and brain (Cechetto and Saper, 1987; Kalia and Mesulam, 1980; Rajendran et al., 2019). In humans, empirical evidence for causal influence in both directions exist, but the findings are considered tentative (Candia-Rivera et al., 2021; De Falco et al., 2024; Pollatos et al., 2005; Sargent et al., 2024; Silvani et al., 2016). Similarly, bi-directional interactions have been observed in other bodily systems, such as respiration (Kluger and Gross, 2021; McKay et al., 2003).

Since our methodological approach did not allow making causal inferences, we cannot specify whether our findings reflect ascending influences of peripheral physiology on neural activity or the cortical regulation of cardiac activity. However, our analysis approach, similar to some earlier studies (Candia-Rivera et al., 2021), was designed to test the modulatory effects of peripheral physiology on neural activity. This experimental design thus allows to make directionality assumptions on the data, even though confirmatory conclusions cannot be made. While these approaches enable noninvasive investigation of heart-brain coupling, development of novel methodological approaches is warranted to comprehensively investigate the complex interplay between afferent and efferent interactions between heart and brain.

In support of the afferent influence, the spatial distribution of the modulatory effects of HRV on oscillatory brain activity observed in this study overlaps with the brain regions previously identified to respond to cardiac activity (Coll et al., 2021; Engelen et al., 2023) and to be involved in parasympathetic regulation, such as temporal and insular cortices (Beissner et al., 2013). The topography is also consistent with the previous studies that have identified extensive neural networks in which different bodily functions govern neural activity (Engelen et al., 2023; Kluger and Gross, 2021; Rebollo and Tallon-Baudry, 2022; Richter et al., 2017). Given that different measures of peripheral physiology (e.g., respiration, electrodermal activity, pupillary response) offer rich and complementary information about the physiological state of the body, their combination (e.g., cardiorespiratory coupling, sympathetic and parasympathetic activity) could in the future enhance our understanding of how various bodily systems contribute to brain functioning and behavior.

Interestingly, the findings of our study and the observations of previous studies have demonstrated that the neural networks modulated by the peripheral physiology overlap with the topography of the well-established functional neural networks, such as the default mode network, dorsal and ventral attention network, salience network, and cingulo-opercular network (Corbetta and Shulman, 2002; Kleckner et al., 2017; Raichle, 2015; Sadaghiani and D’Esposito, 2015; Seeley, 2019). These brain regions are involved both in monitoring and regulating the peripheral physiology, and in a variety of psychological processes including attention, emotions, social information processing, and decision-making (Beissner et al., 2013; Corbetta and Shulman, 2002; Kleckner et al., 2017; Sklerov et al., 2019; Thayer et al., 2012; Van Overwalle, 2009). This result hints at the idea that interaction between peripheral physiology and neural activity may be more intertwined with subjective experience and behavior than previously thought.

Our findings add to the existing research on heart-brain coupling by focusing on derived measures of cardiac signaling, such as HRV utilized in this study, that were recently recognized as bringing potentially more comprehensive understanding of heart-brain coupling (Godwin et al., 2024; Candia-Rivera et al., 2024; Parviainen et al., 2023). We also evidenced that derived measures provide a feasible metric to approach the relevance of heart-brain coupling for individual trait characteristics. Furthermore, indicated by the present and earlier findings by our research group (Heinilä et al., 2024), it may be that a well-defined contrast between conditions or tasks more readily induces brain activity that is relevant for individual trait characteristics.

In accordance with our first hypothesis, we demonstrated that the fluctuations in heart-brain coupling are associated with individual trait characteristics, namely behavioral approach and avoidance tendencies. While a smaller modulatory effect of HRV on alpha power in left temporoparietal cluster was associated with stronger behavioral avoidance tendency, a larger modulatory effect of HRV on beta power in left occipital and temporal clusters was associated with stronger behavioral approach tendencies, BAS drive and BAS fun seeking, respectively. Together our findings indicate that individuals with stronger interaction between cardiac and neural signaling report to be less prone to avoid negative outcomes, tend to seek novel experiences, and are committed to pursuing desired goals.

Surprisingly, even though the association between temperament and physiological reactivity is well-established (Kagan et al., 1987; Knyazev et al., 2002), only individual studies have examined the relationship between body-brain interaction and individual trait characteristics. Shokri-Kojori et al. (2018) observed that the degree of synchrony between peripheral signals and neural activity is associated with temperamental characteristics. This observation is in accordance with our findings and further highlights the importance of understanding the role of body-brain interaction in characterizing temperament and personality.

Our study aligns with earlier evidence on the relevance of temporoparietal oscillatory brain activity for individual trait characteristics (Coan and Allen, 2003; Heinilä et al., 2024; Kennis et al., 2013; Knyazev et al., 2002; Smith et al., 2017; Sutton and Davidson, 1997). However, there are also conflicting findings (Coan and Allen, 2003; Schneider et al., 2016) and the inconsistencies in the literature may be attributed to the high variability in the experimental settings or methods used. Especially, asymmetry measures widely used in this field have sparked some controversy (Smith et al., 2017; Vecchio and De Pascalis, 2020) and may explain at least some of the inconsistent findings. Moreover, while self-report questionnaires provide a valuable method to assess individual trait characteristics, they reflect the potential neural and physiological correlates of temperament only indirectly and may raise concerns regarding constraints of self-awareness and response bias (Chan, 2010; Paulhus and Vazire, 2007).

In addition, or alternatively, to body-brain interaction itself, also the attunement to internal bodily sensations may be a critical factor in influencing individual trait characteristics. This was suggested by the findings of our moderator analysis, where the attunement to internal bodily sensations moderated the relationship between heart-brain coupling and behavioral tendencies. However, this effect was only observed for the association between beta power modulation and behavioral approach tendency, and not for the link between alpha power modulation and behavioral avoidance tendency. Specifically, a positive relationship between heart-brain coupling in the beta band and behavioral approach tendency was observed only in individuals with a stronger attunement to visceral information.

Thus, our findings support the hypothesis that the association between heart-brain coupling and behavioral tendencies is at least partly moderated by the attunement to internal bodily sensations. This finding corroborates the previous findings indicating that the stronger tendency to attend to the internal milieu could predispose individuals to greater awareness of the bodily physiology and potentially also amplify the bodily sensations and responsiveness through the attunement to the internal milieu (Lyyra and Parviainen, 2018). While still rather tentative, these findings emphasize the need for revisiting the neurobiologically motivated theoretical accounts of temperament to also consider body-brain interaction and attunement to internal bodily sensations in understanding individual trait characteristics. Further studies are essential to gain deeper insights into the association between body-brain interaction and individual trait characteristics across the lifespan. Especially, the use of long-term measurements (days to weeks), longitudinal experimental designs, and ecologically valid research settings could significantly improve our understanding of the extent to which body-brain interaction and attunement to internal milieu shape individual trait characteristics.

## Conclusions

This work contributes to the existing knowledge of modulatory effects of peripheral physiology on neural activity by demonstrating alpha and beta power increases during states of lower parasympathetic activity. Our findings also bring insights to the intricate interplay between cardiac and neural signaling and its relation to individual trait characteristics. Specifically, we showed that heart-brain coupling was associated with behavioral approach and avoidance tendencies and that the relationship between heart-brain coupling and individual trait characteristics is at least partly moderated by the attunement to internal bodily sensations. These findings have significant implications for the understanding of how bodily functions govern the ongoing oscillatory brain activity across the cortex. Moreover, our findings demonstrate that not heart-brain coupling as such but its moderation by the attunement to the bodily sensations contributes to the way individual approaches and interacts with the external world.

## Acknowledgements

This work was supported by the Alfred Kordelin Foundation, the Finnish Cultural Foundation, and the Central Finland Regional Fund of the Finnish Cultural Foundation. The funders had no role in study design, data collection and analysis, decision to publish, or preparation of the manuscript. We wish to express our thanks to Hanna Honkanen, Hanna-Maija Lapinkero, Sannamari Matveinen, Emilia Penttinen, and Reetta Siikavirta for their assistance during data collection. We are also grateful to Joona Muotka for the help with the statistical analysis.

## Data availability

The data cannot be made openly available according to the ethical permission and national privacy regulations at the time of the study, but the data supporting the findings of this study are available upon reasonable request from the authors and with permission of the Ethics Committee of the University of Jyväskylä.

## Contributions

SK: Conceptualization, Methodology, Formal analysis, Investigation, Writing – Original Draft, Writing – Review & Editing, Funding acquisition

JK: Conceptualization, Methodology, Formal Analysis, Software, Writing – Review & Editing

TA: Conceptualization, Writing – Review & Editing, Supervision

TP: Conceptualization, Methodology, Formal analysis, Writing – Review & Editing, Supervision

## Appendix

**Fig. A.1.**
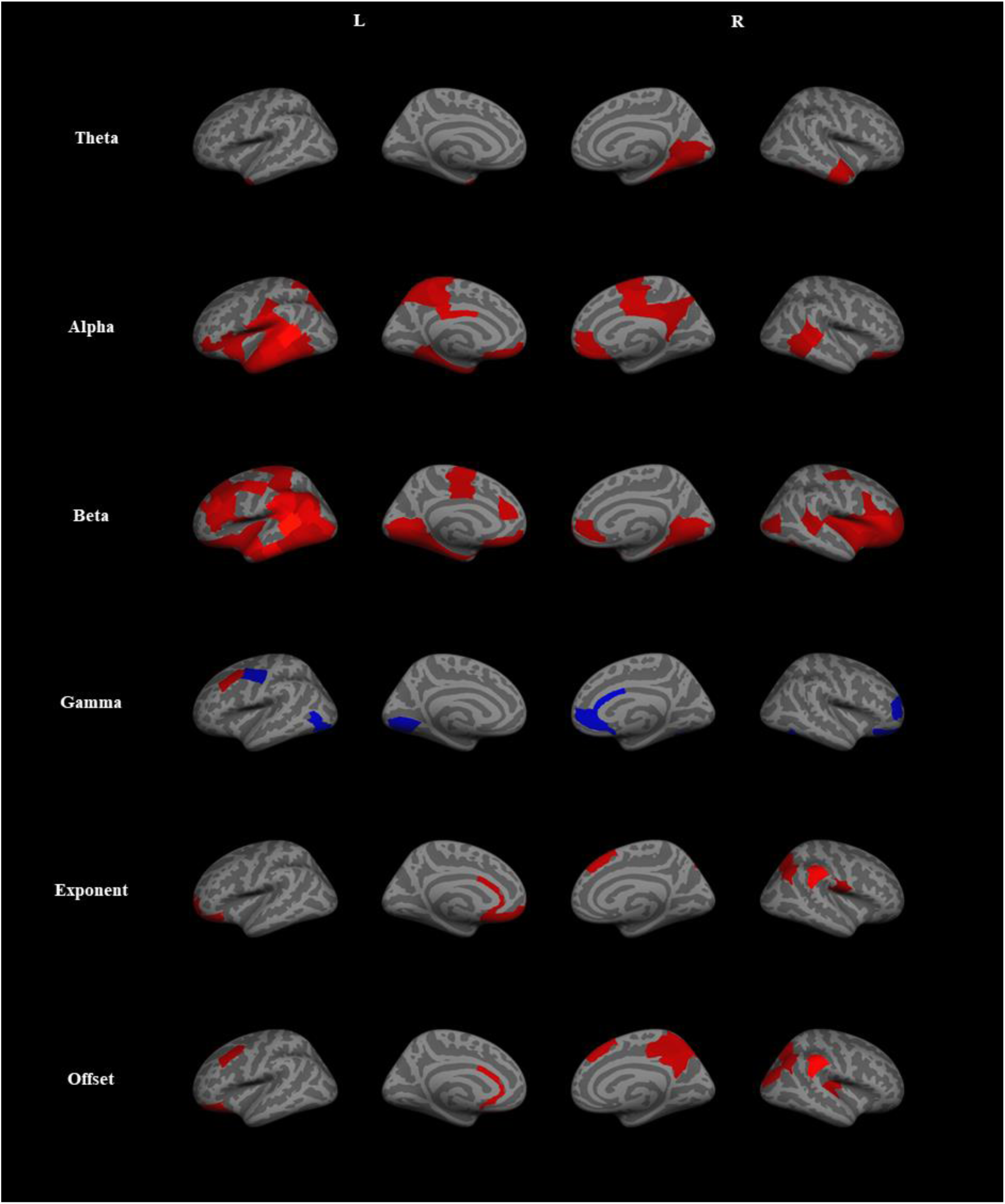
Modulations of periodic and aperiodic activity between low vs. high HRV states. Statistically significant differences (*p* < 0.05, uncorrected) were observed in periodic (theta, alpha, beta, and gamma power) and aperiodic activity (exponent and offset) between low vs. high HRV states. Blue color indicates statistically significant decreases and red color statistically significant increases in periodic and aperiodic activity between low vs. high HRV states.

